# Dinoflagellate Proton-Pump Rhodopsin Gene in Long Island Sound: Diversity and Spatiotemporal Distribution

**DOI:** 10.1101/2022.08.24.505117

**Authors:** Huan Zhang, Kelly J. Nulick, Zair Burris, Melissa Pierce, Minglei Ma, Senjie Lin

## Abstract

Microbial proton-pump rhodopsin (PPR), an alternative light-harvesting mechanism to chlorophyll-based photosystems, may contribute significantly to solar energy entry into the marine ecosystem. PPR transforms solar energy to cellular energy used for various metabolic processes in the cells or flagellar movement. Although rhodopsins or their encoding genes have been documented in a wide phylogenetic range of cultured dinoflagellates, information is limited about how widespread and how spatiotemporally dynamical dinoflagellate PPR (DiPPR) are in the marine ecosystem. In this study, using the quantitative PCR method, we investigated the abundance and diversity of DiPPR genes in Long Island Sound (LIS), a temperate estuary, both spatially and temporally in 2010. DiPPR genes were found year-round and throughout LIS, with higher abundances in the eutrophic Western Sound and during April and July. The gene diversity data suggest that there are at least five distinct rhodopsin-harboring groups of dinoflagellates throughout the year. The abundance of DiPPR genes, measured as copy number per mL seawater, appeared not to be influenced by water temperature and nitrogen nutrient concentration; however, weak negative correlations with orthophosphate concentration and salinity and a positive correlation with the abundance of typical-DiPPR-harboring dinoflagellates were observed. The association of DiPPR with phosphorus nutrition warrants further studies.

## Introduction

Virtually all forms of life on the surface of the earth are driven by sunlight energy. Prior to the year 2000, chlorophyll-based photosynthesis had been considered the sole mechanism by which aquatic organisms draw energy from sunlight. Marine phytoplankton contribute about 50% of global photosynthetic productivity [1,2]. However, with the discovery of proton-pump rhodopsin (PPR) in marine bacteria in 2000 [3,4], it has become clear that rhodopsin-based phototrophy is responsible for capturing a substantial portion of the solar energy that enters the marine ecosystem daily [5–7]. PPRs are now found in various archaea, bacteria, fungi, and algae (e.g., [3,4,8–11]). With few exceptions, PPRs transform solar energy to cellular energy, which can support various cellular functions, such as ATP synthesis, substrate transportation, and survival of bacteria under carbon starvation [6,9]. Metagenomics data show that 13-60% of the bacterial genomes in surface ocean possess PPR genes [12–14]. Since each rhodopsin binds a single molecule of retinal, the total number of retinal molecules is equivalent to that of rhodopsins. Gómez-Consarnau et al. [7] quantified the vertical distributions of the *all-trans* retinal and two other energy-converting pigments, chlorophyll-*a,* and bacteriochlorophyll-*a*, along a nutrient gradient in the Mediterranean Sea and the Atlantic Ocean. They discovered that PPR-based phototrophy likely contributes the same amount of light energy fixation as chlorophyll-*a* based phototrophy does, and that the energy obtained by PPRs is sufficient to support basal metabolism of bacteria in the surface ocean. Their study provides quantitative evidence that PPR phototrophy is a major light energy harvesting mechanism in the surface ocean.

Among the rhodopsin-harboring microbial eukaryotes, PPR genes seem to be the most widespread in dinoflagellates [10]. Initially detected in *Pyrocystis lunula* through microarray analysis [15], later through transcriptomic studies on lab cultures, PPR genes have been documented in basal lineages such as the heterotrophic lineage *Oxyrrhis marina* [10,16,17], autotrophic lineages *Polarella glacialis* [8], *Prorocentrum donghaiense* [18], the mixotrophic lineage *Karlodinium veneficum* [19, this study], to the evolutionarily most recent lineage *Alexandrium* [10]. Except those from *P. shikokuense* (formerly *P*. *donghaiense)* and *K. veneficum,* most of the dinoflagellate PPR genes (referred to as typical dinoflagellate PPR, or DiPPR hereafter), share high similarity to PPRs from the gamma-proteobacteria of the SAR86 group [3]. Although we previously detected two DiPPR cDNAs from a metatranscriptome of a natural plankton assembly [8], thus far, no studies have been dedicated to investigating the spatial and seasonal variation of DiPPR in the field, although some data generated from dinoflagellate blooms [20,31], diatom-dominated community [11], or general microbial eukaryotes [5] are available. In this study, we isolated six novel full-length DiPPR cDNAs from a boat dock near the Avery Point campus of the University of Connecticut in the eastern section of Long Island Sound (LIS). Furthermore, we designed primers on the conserved regions of the cDNAs and used quantitative PCR to analyze the spatial and temporal variations of the abundance and diversity of DiPPR genes over the course of one year from the eutrophic western end to the mesotrophic eastern end of the Sound.

## Methods

### Water sample collection

Water samples were collected by the personnel of the Long Island Sound (LIS) Water Quality Monitoring Program from the Connecticut Department of Energy and Environmental Protection. Surface water sampling was carried out each month from January to December 2010, from ten stations (A4, B3, C1, D3, E1, F2, H4, I2, J2, and K2) except when inclement weather prevented cruises (Fig. 1). Two hundred ml of water samples were collected 2 m below the water surface. For preservation of phytoplankton cells, four mL of neutral Lugol’s (Utermöhl’s) solution were added to each water sample (final concentration 2%) as reported [22].

**Fig. 1.**
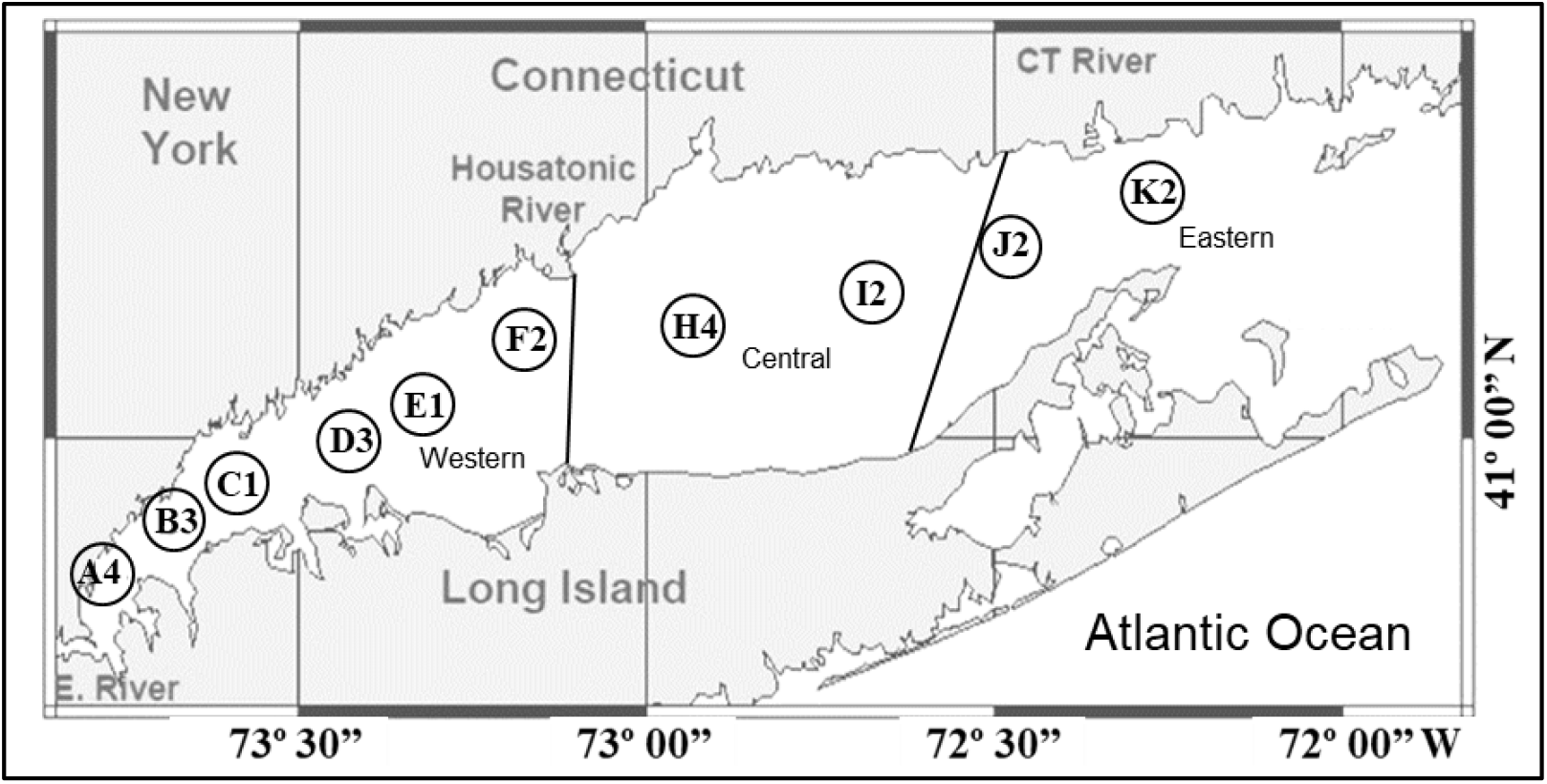
Sampling stations in Long Island Sound.

A total of 95 water samples were collected (Table 1). The preserved samples were kept at 4°C in darkness until processing generally within 2 months. For each sample, a 50mL-subsample was concentrated using Utermöhl Settling Chamber for at least 48 hours. The supernatant was aspirated so that the sample was concentrated to 1mL and examined using an Olympus BX51 microscope (Olympus) equipped with 10 and 40 x magnification on the objective and 10 x in the eyepiece to achieve up to 400 x magnification. Phytoplankton were identified to the lowest taxonomic level possible, usually to the genus level. A parallel set of the samples were taken for genomic DNA (gDNA) extraction.

**Table 1.**
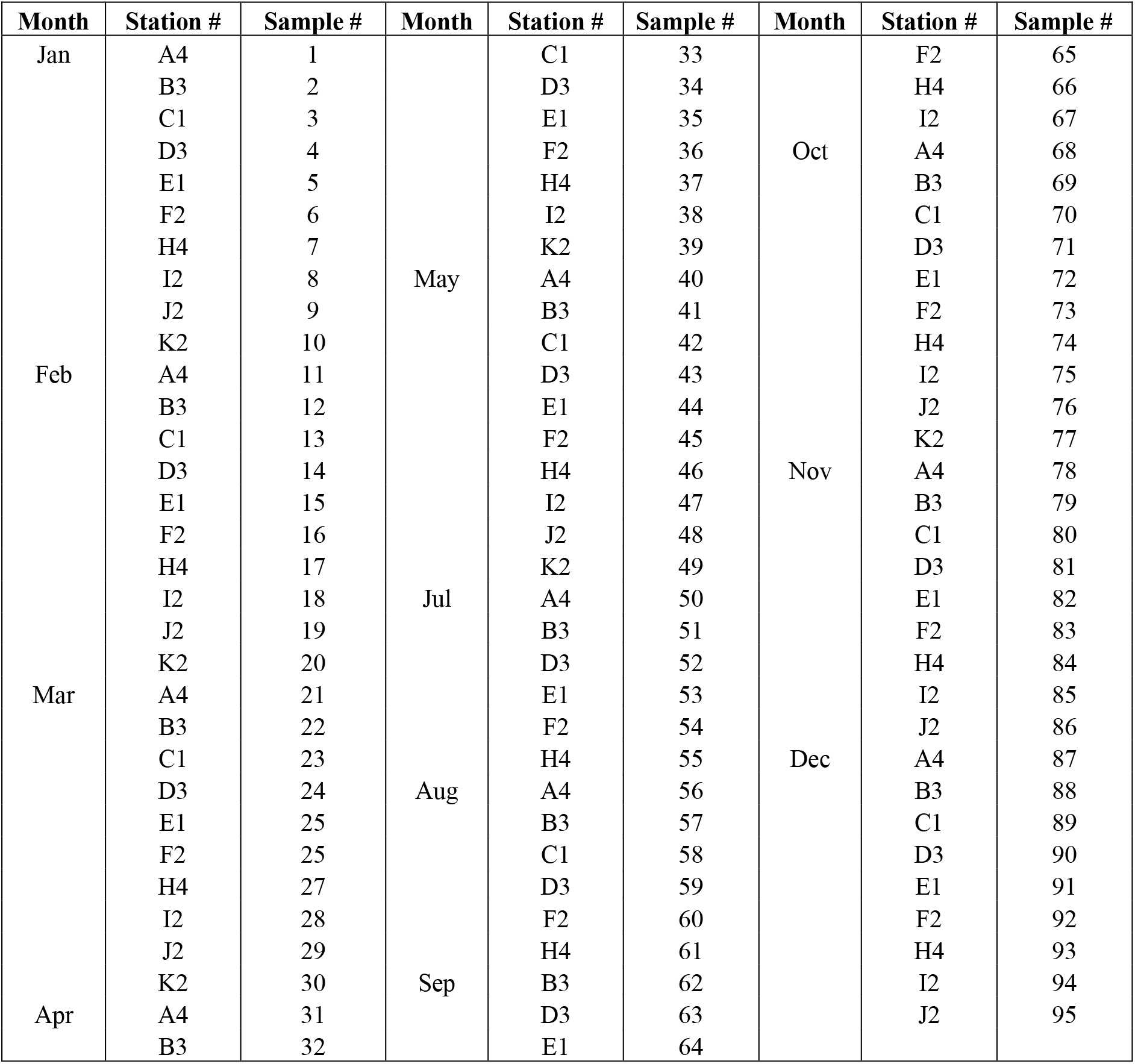
Stations and months of water samples collected in this study.

### Genomic DNA extraction and PCR amplification of 18S rRNA and rhodopsin genes

Genomic DNA was extracted from LIS water samples according to [23] with some modification. Briefly, for each water sample, a 50mL-subsample was concentrated to 1mL as mentioned above, transferred to a 2mL microtube, and centrifuged at 15,000 g for 2 min to pellet the cells. After the supernatant was removed carefully, approximately 100 mg of 0.5 mm-zirconia/silica beads (Biospec Products) were added to the pellet and bead-beaten at 6m/s for 30 seconds using an MP Fast Prep-24 Tissue and Cell Homogenizer (MP Biomedicals).

250 μL CTAB buffer (2% Cetyl Trimethyl Ammonium Bromide, 100mM Tris-HCl pH8, 20mM EDTA pH8, 1.4M NaCl, 0.2% β-mercaptoethanol, 0.1 mg/ml Proteinase K) was added, and the samples were incubated for 4 hrs at 55 °C with gentle mixing. For each sample, 250μL chloroform was added and mixed well, centrifuged for 10 minutes at 15, 000g, and the supernatants transferred to a new 2mL microtube. The genomic DNA was then purified using DNA Clean & Concentrator Kit (Zymo Research) and eluted in 200 μl of 10mM Tris-HCl (pH8), making the final amount of DNA per 4 μL equivalent to the DNA from 1 mL water sample. DNA quantity and quality were determined using NanoDrop Spectrophotometer ND-1000 (NanoDrop Technologies) and stored at −20 °C until further analysis.

### Development of specific primers for typical dinoflagellate proton-pump rhodopsin (DiPPR) genes and quantitative Real-Time quantitative PCR (qPCR)

Based on the alignment of the sequences of typical DiPPR sequences available in GenBank databases, several conserved regions were identified, and primers were designed in these regions using Primer Premier 6.0 (Table 2). These primers were tested against the cDNAs/ genomic DNAs (gDNAs) of the Phytoplankters we obtained in the previous study (see [24] for details). Many of these phytoplankton species were commonly found in LIS (Table 3; http://media.ctseagrant.uconn.edu/publications/marineed/phytoplankton/phytoplankton.pdf).

**Table 2.**
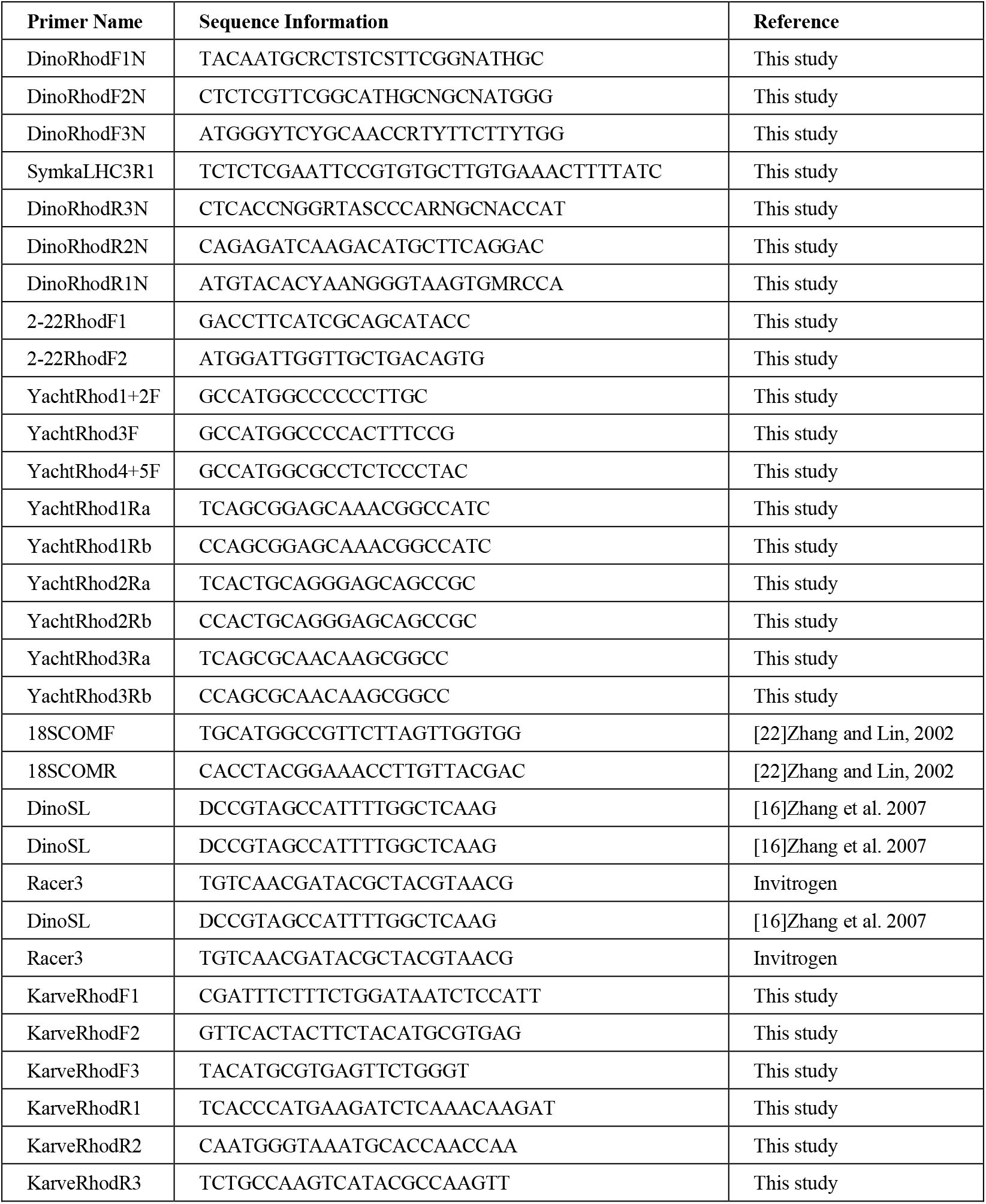
Primers used in the present study.

**Table 3.** Phytoplankton species tested for dinoflagellate rhodopsin primers.

| Primer name | Strain | cDNA | gDNA |
| --- | --- | --- | --- |
| *Alexandrium catenella* (formally *A. fundyense*) | CCMP1719 | + | + |
| *Alexandrium pacificum* | ACHK | + | - |
| *Amphidinium carterae* | AC | + | + |
| *Ditylum brightwellii* | CCMP 2227 | - | - |
| *Dunaliella tertiolecta* | CCMP1320 | - | - |
| *Emiliania huxleyi* |  | - | - |
| *Heterocapsa triquetra* | CCMP 448 | - | - |
| *Heterosigma akashiwo* | CCMP 452 | - | - |
| *Karenia brevis* | Wilson | - | - |
| *Karlodinium veneficum* | CCMP1975 | - | - |
| *Odontella sinensis* | CCMP 1815 | - | - |
| *Oxyrrhis marina* | CCMP1795 | + | + |
| *Pfiesteria piscicida* | CCMP1831 | - | - |
| *Pfiesteria shumwayae* | CCMP2359 | - | - |
| *Polarella glacialis* | CCMP2088 | + | + |
| *Prorocentrum donghaiense* |  | - | - |
| *Prorocentrum minimum* | CCMP696 | - | - |
| *Pyrocystis lunula* |  | + | - |
| *Pyrocystis noctiluca* | CCMP732 | + | - |
| *Rhodomonas* sp. | CCMP768 | - | - |
| *Skeletonema costatum* | China30 | - | - |
| *Fugacium kawagutii* (formerly *Symbiodinium kawagutii*) | CCMP2468 | - | - |
| *Symbiodinium microadriaticum* | CCMP 2458 | + | - |
| *Symbiodiniaceae* sp. | SSBO1 | - | - |
| *Thalassiosira pseudonana* | CCMP1335 | - | - |

Primer sets DinoRhodF2N-DinoRhodR2N and DinoRhodF3N-DinoRhodR3N were found to be sensitive and specific to amplify rhodopsin gene for many dinoflagellates (Table 3). To assess spatial and temporal dynamics of rhodopsin gene abundance, Real-Time qPCR was performed as reported [25] using 4 μL of the extracted DNA as the template with primer set DinoRhodF3N-DinoRhodR3N. A dilution series of the PCR product of *Alexandrium catenella* (formerly *A. fundyence)* rhodopsin cDNA was used as a standard to calculate the copy number of this gene in LIS DNA samples as reported [23].

As a reference with which to normalize rhodopsin gene abundance in phytoplankton communities, we also performed Real-Time qPCR for the 18S rRNA gene (18S rDNA) in all 95 samples using the previously reported universal primer set 18ScomF-18ScomR [23]. This primer set has been verified to be able to amplify all the major phytoplankton.

#### DiPPR gene cloning and sequencing of the LIS water samples

Six DNA samples were chosen for cloning and sequencing of the DiPPR genes, including three samples from station A4 (during February, April, and August, sample # 11, 31, and 56, respectively), one sample from station E1 in April (sample # 35), and two samples from station H4 (February and August, sample # 7 and 61). To measure rhodopsin diversity in these samples, amplicons from the 2^nd^ round of rhodopsin Real-Time qPCR were cloned into T-vectors, following our previous reported method [22]. For each sample, 48 colonies were randomly chosen, plasmid DNA isolated, and sequenced as reported [22].

### RNA extraction, cDNA synthesis and full-length cDNA isolation of DiPPR in the field and in *Karlodinium veneficum* strain CCMP1975

Water samples were collected in February 2011 at the Yacht Club marina (41°18’55”N, 72°O3’48.6”’W) at the Avery Point campus of the University of Connecticut. One liter of water was filtered onto a 3-μm Nuclepore filter membrane which was then immediately immersed in 1 mL TRIzol RNA buffer. In parallel, ~1 million cells of *Karlodinium veneficum* CCMP1975 were harvested from a culture grown on f/2 medium, by centrifuging at 3000g, and fixing cell pellet in 1 mL TRIzol RNA buffer. Total RNA was extracted following the reported method [16]. First-strand cDNA was synthesized and purified, full-length rhodopsin cDNAs were obtained following the reported DinoSL-based method [16] using the DiPPR primers pared with DinoSL or Racer3 primer (Table 2).

### Phylogenetic analysis

All the dinoflagellate rhodopsin sequences reported in GenBank were obtained by BLAST search. The rhodopsin nucleotide sequences we obtaiend in this study and the representatives acquainted from the GenBank database were aligned using ClustalW in MEGA X. Model in MEGA X were used to find the best model of Nuclear acid evolution. Phylogenetic trees were inferred using Maximum likelihood method in MEGA X with rates estimated from Model in MEGA X. The evolutionary history was inferred by using the Maximum Likelihood method with Hasegawa-Kishino-Yano model [26]. The bootstrap consensus tree inferred from 1000 replicates [27] is taken to represent the evolutionary history of the taxa analyzed [27]. Branches corresponding to partitions reproduced in less than 50% bootstrap replicates are collapsed. The percentage of replicate trees in which the associated taxa clustered together in the bootstrap test (1000 replicates) are shown next to the branches [27]. Initial tree(s) for the heuristic search were obtained automatically by applying Neighbor-Join and BioNJ algorithms to a matrix of pairwise distances estimated using the Maximum Composite Likelihood (MCL) approach, and then selecting the topology with superior log likelihood value. A discrete Gamma distribution was used to model evolutionary rate differences among sites (5 categories (+G, parameter = 0.4893)). This analysis involved 202 nucleotide sequences. All positions containing gaps and missing data were eliminated (complete deletion option). There were a total of 230 positions in the final dataset. Evolutionary analyses were conducted in MEGA X [28]. The resulting tree file from MEGA X was then uploaded to the Evolview to make further modifications.

### Statistical analyses of DiPPR/18S rDNA gene abundance and relationships with environmental factors

To examine the potential correlation between the DiPPR/18S rDNA gene abundance and environmental factors, we obtained the monthly data for the 10 stations including salinity, water temperature, chlorophyll *a*, concentration of ammonia, nitrate + nitrite, orthophosphate, total dissolved nitrogen and total dissolved phosphorus in 2010 (http://lisicos.uconn.edu/dep_portal.php, Fig. 2). The statistical analyses of the relationship between environmental factors and the corresponding gene abundances of DiPPR and 18S rRNA in each sample were carried out using the built-in Regression function in Microsoft Office Excel 2016 Data Analysis.

**Fig. 2.**
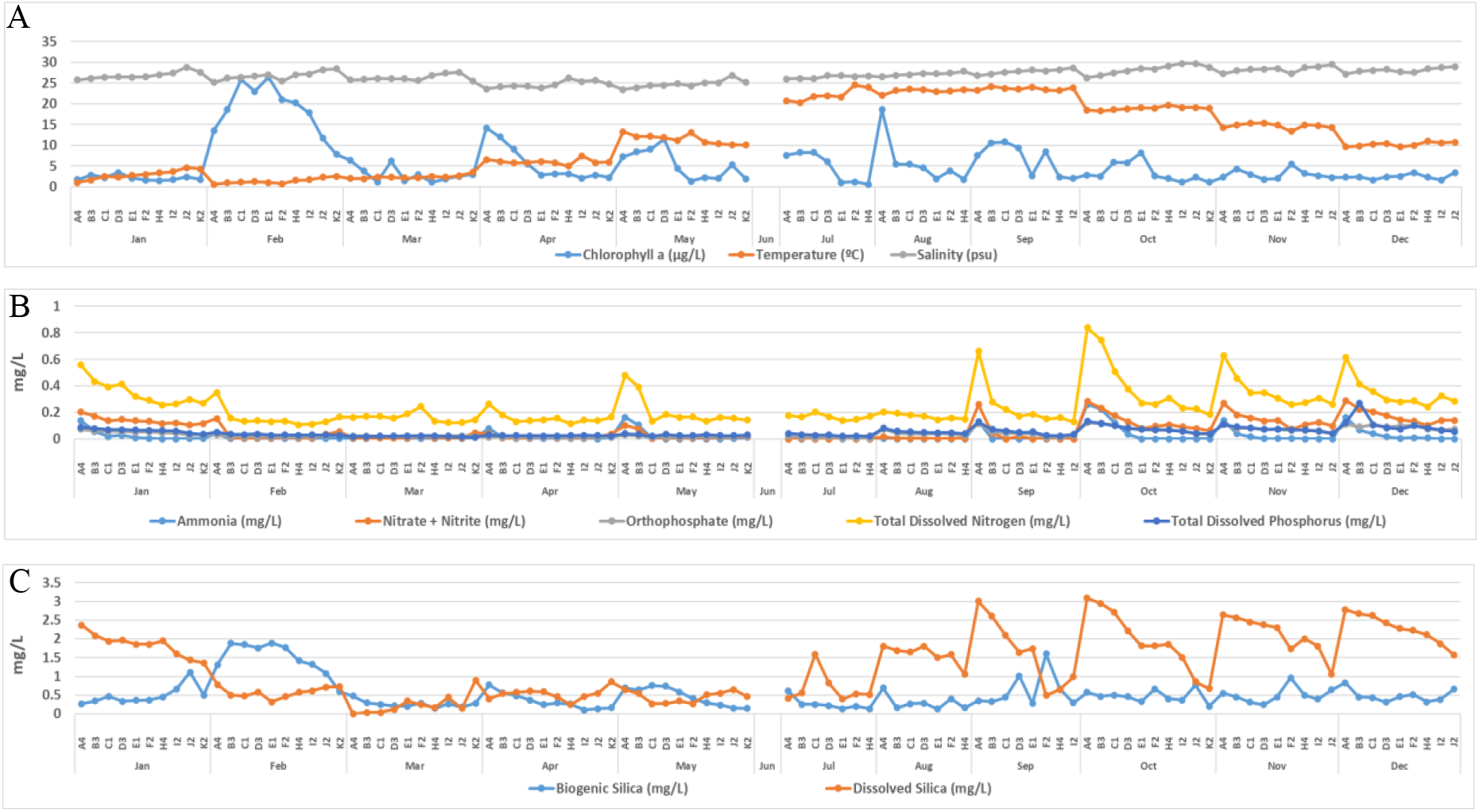
Monthly change of environmental factors in 2010, Long Island Sound. A) Monthly changes of Chlorophyll *a,* temperature and salinity. B) monthly changes of concentration of ammonium, nitrate + nitrite, orthophosphate, total dissolved nitrogen and total dissolved phosphorus in 2010 Long Island Sound. C) monthly changes of biogenic silica and dissolved silica concentration in 2010, Long Island Sound.

## Results

### Temporal and spatial variations of phytoplankton cell abundance

The phytoplankton community showed large temporal and spatial changes (Fig. 3, Fig. S1A). The average cell abundance showed two peaks, one in February and the other in September, with cell concentrations of 22.09±9.45 x10^5^ and 26.26±23.60 x10^5^ cells•L^-1^, respectively. The lowest cell abundance was found in April, which was 3.49±1.58 x10^5^ cells•L^-1^. Diatoms dominated throughout the whole year, while dinoflagellate cell concentration was highest in July, with the most abundant species from the genera *Gymnodinium, Amphidinium, Heterocapsa, Prorocentrum,* and *Scrippsiella.* The other phytoplankton, including silicoflagellates, cryptophytes, ochrophytes, haptophytes, euglenozoa, and unidentified small flagellates and nanoplankton reached minor peaks in May and December (Fig. 3A, Table S1).

**Fig. 3.**
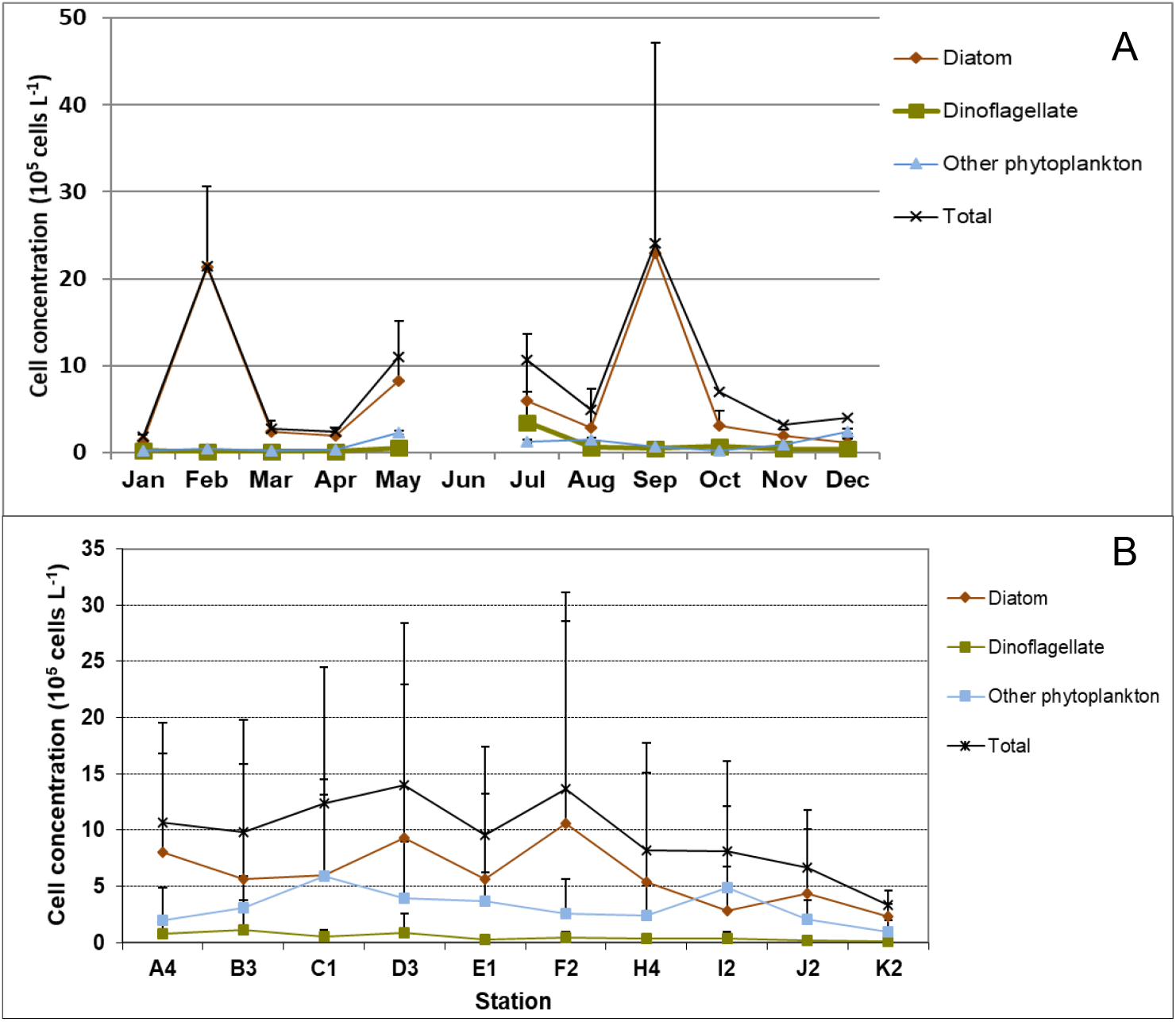
Temporal and spatial dynamics of Long Island Sound (LIS) phytoplankton in 2010. A) Monthly average of phytoplankton cell concentration of diatom, dinoflagellate, other phytoplankton and total phytoplankton for all stations in LIS. B) Annual average of phytoplankton cell concentration of diatom, dinoflagellate, other phytoplankton and total phytoplankton in each station.

Annual average phytoplankton abundance was generally higher in the western LIS than the eastern Sound. Overall, cell concentrations were higher at stations A4-F2 than stations east of them (H4-K2). The highest level of abundance appeared to be in D3 (14.02±14.37 x10^5^ cells•L^-1^), although the difference was not statistically significant (p>0.05) due to large seasonal variations. Annual average cell concentrations for stations in the western sound (A4 to F2) were 9.6-14.02 x10^5^ cells•L^-1^, those in the central sound (H4, I2) were 8.1-8.2 x10^5^ cells•L^-1^, and those in the eastern sound (J2, K2) were 3.3-6.6 x10^5^ cells•L^-1^(Fig. 3B).

### Seasonal changes of gene abundance of DiPPR and phytoplankton 18S rDNA in LIS

Genomic DNA was isolated successfully from all 95 water samples obtained and used for RealTime qPCR. The DiPPR gene was detected at every station with abundance ranging from several copies to up to 1.3 million copies•mL^-1^ (Fig. 4A, Fig. S2B), and the average monthly abundance showed two major peaks in April and July; however, when DiPPR gene abundance was normalized to the monthly average dinoflagellate cell number, only the April peak remained (4.18 x10^4^ copies/cell; Fig. 4B). The total phytoplankton18S rDNA abundance ranging from 265 thousand copies to over 36.9 million copies•mL^-1^, with a major peak in February and two minor peaks in July and September, respectively, similar to the pattern of the seasonal change of the total phytoplankton cell abundance (Figs. 3, 4).

**Fig. 4.**
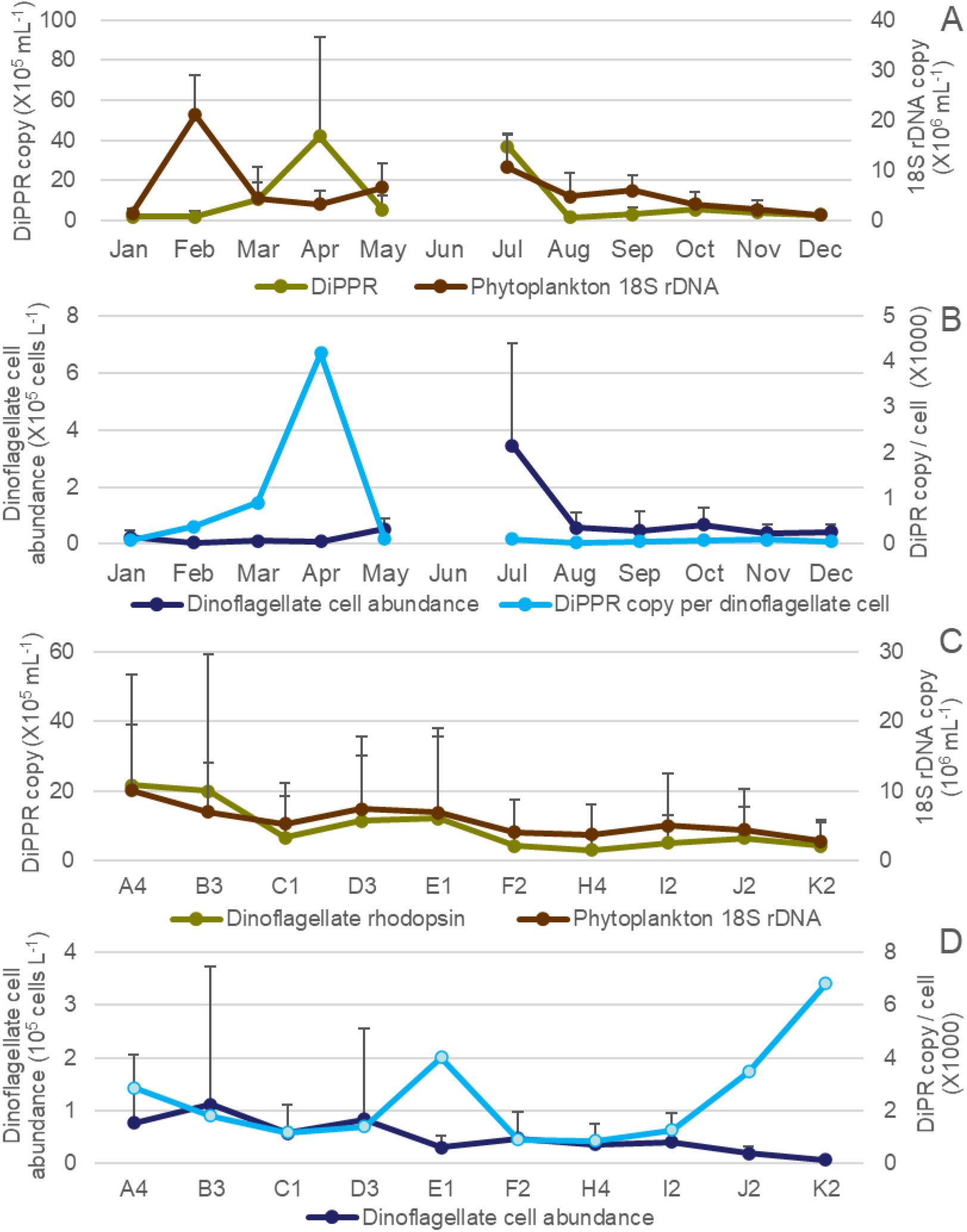
Temporal and spatial profiles of dinoflagellate proton-pump rhodopsin (DiPPR) and phytoplankton 18S rDNA gene abundance in 2010. LIS. A. temporal profile of average DiPPR and phytoplankton 18S rDNA gene abundance; B. temporal profile of average dinoflagellate cell abundance and DiPPR copy per cell; C, spatial profiles of average DiPPR and phytoplankton 18S rDNA gene abundance; D. spatial changes of average dinoflagellate cell abundance and DiPPR copy per cell.

The spatial change of both DiPPR and phytoplankton 18S rDNA gene abundances showed a weak decreasing pattern from the western sound to the central and eastern sound, with large variation within each station in each month (Fig. 4C, Fig. S1B, S1C). This pattern was similar to that of phytoplankton cell abundance (Fig. 3) in LIS. However, DiPPR gene abundance showed no significant correlations with temperature, salinity, nutrient concentration and chlorophyll *a* (Figs. S2–S4).

When DiPPR gene abundance was normalized to the monthly average dinoflagellate cell abundance, peaks appeared at stations E1 and J2-K2 (4.00 x10^3^, 3.46 x10^3^, 6.82 x10^3^ copies/cell, respectively; Fig. 4D). The highest levels of DiPPR and phytoplankton 18S rDNA gene abundance were both at station A4 (2.17±3.18 x10^5^ copies•mL^-1^ and 10.07±9.55 x10^6^ copies•mL^-1^, respectively), with the lowest at station H4 (0.30±0.36 x10^5^ copies•mL^-1^ and 3.72±4.35 x10^6^ copies/mL^-1^, respectively; Fig. 4C, Fig. S1B). Annual average gene abundances were 4.2-21.7 x10^4^ copies•mL^-1^ and 4.03-10.07 x10^6^ copies•mL^-1^ in western sound stations (A4 to F2), 3.0-4.9 x10^4^ copies•mL^-1^ and 3.72-5.03 x10^6^ copies•mL^-1^ in the central sound (H4, I2), and 4.1-6.4 x10^4^ copies•mL^-1^ and 2.80-4.36 x10^6^ copies•mL^-1^ in the eastern sound (J2, K2), respectively for DiPPR and phytoplankton 18S rDNA (Fig. 4).

### Correlation between DiPPR/18S rDNA gene abundance and environmental factors

The relationship of DiPPR and 18S rRNA gene abundances with environmental factors are shown in Figs. S2–S4. DiPPR gene abundance was not clearly correlated with concentration of ambient nitrogen-nutrient (ammonia, nitrate+ nitrite, and total dissolved nitrogen), dissolved phosphorus, chlorophyll *a*, or ambient water temperature. However, DiPPR copy number showed a weak negative linear correlation with orthophosphate concentration (two-way ANOVA df=1, F=5.896; R^2^ = 0.060, p< 0.05) and salinity (df=1, F=15.769; R^2^ = 0.145, p< 0.01). On the other hand, Phytoplankton18S rDNA gene abundance was not clearly correlated with ammonia concentration, salinity or ambient water temperature; however, a strong linear positive relationship with chlorophyll *a* concentration (df=1, F= 164.187; R^2^ = 0.638, p<0.01), and a weak negative correlation with the concentration of nitrate+ nitrite (df=1, F= 10.135; R^2^ =0.098, p< 0.01), total dissolved nitrogen (df=1, F= 6.354; R^2^ = 0.0640, p< 0.05), orthophosphate (df=1, F= 11.994; R^2^ = 0.114, p< 0.01), and dissolved phosphorus (df=1, F= 6.595; R^2^ =0.0662, p< 0.05) was demonstrated (Figs. S2–S4).

**Fig. S1.**
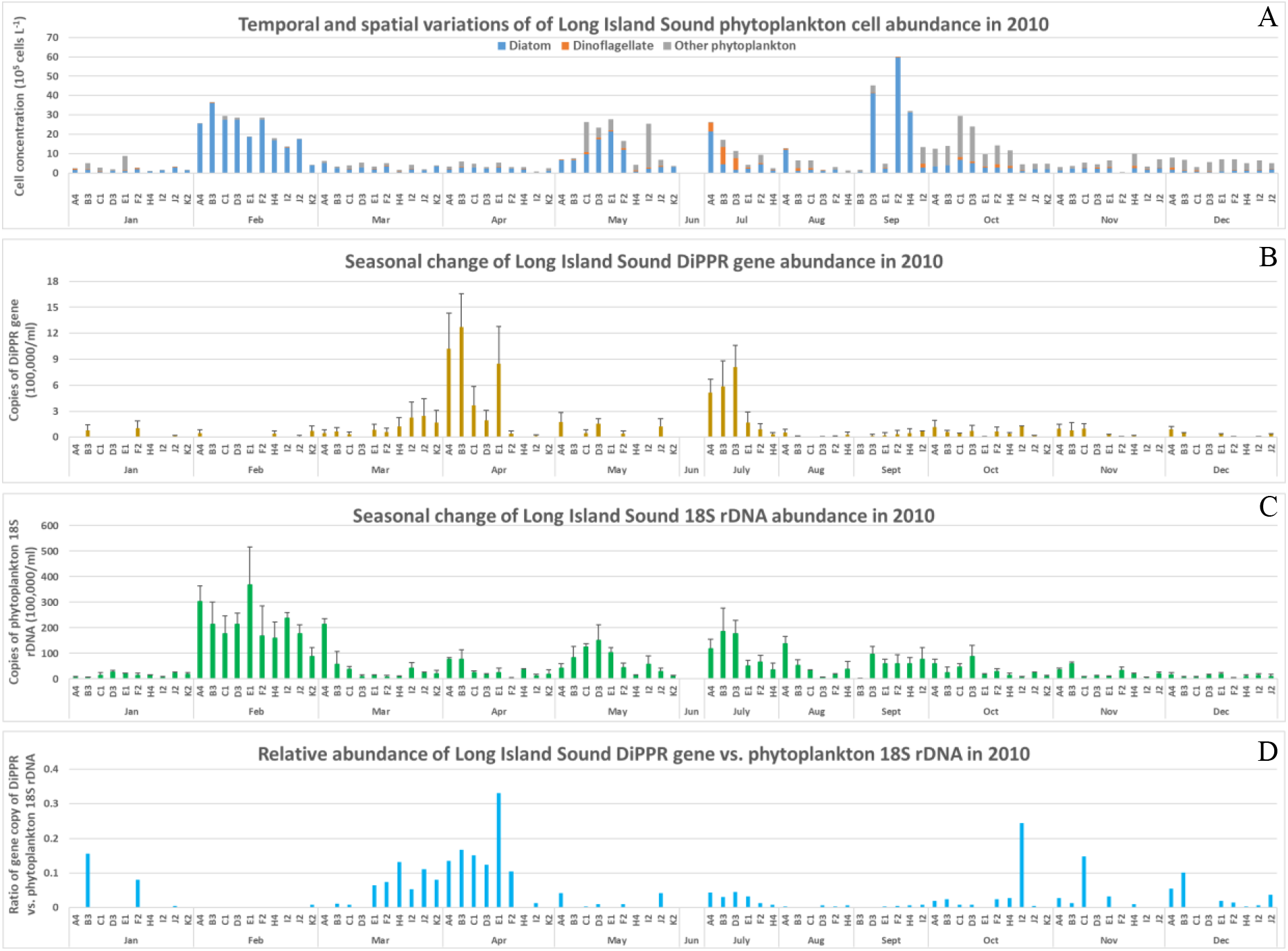
Seasonal changes of phytoplankton cell abundance (A), dinoflagellate rhodopsin (B), phytoplankton 18S rDNA abundance (C) and relative ratio of gene abundance of dinoflagellate rhodopsin vs. phytoplankton 18S rDNA (D) in 2010, LIS.

### Correlation between DiPPR/18S rDNA gene abundance and total phytoplankton cell abundance

The correlation of DiPPR/18S rRNA gene abundances with total community phytoplankton cell abundance is shown in Fig. S4I. DiPPR gene abundance did not show clear correlation with community cell abundance, however, phytoplankton18S rDNA copy number showed a positive correlation with total community phytoplankton cell abundance (df=1, F=39.039; R^2^ = 0.295, p< 0.05).

### DiPPR gene diversity in LIS

Two hundred and nine sequences were obtained from the genomic DNA isolated from six water samples, i.e., A4 in February, April, and August, E1 in April, and H4 in February and August (Table S2). These sequences were used in the diversity analysis along with the representatives of the reported dinoflagellate rhodopsin sequences as well as the six full-length rhodopsin sequences obtained in this study (YachtDinoRhod1, 2, 3, 4, 5 and 6; Fig. 5). Most of the sequences were 306 bp long excluding the primers; however, in most A4 February clones (19 out of 24), there was a 2-nt insertion in the sequence, resulting in frameshift; therefore, these sequences are considered rhodopsin pseudogenes. Eighty-five (41%) of the sequences had unique putative amino acid sequences, indicating that the DiPPR gene is diverse in LIS. The most diverse samples were from A4 and E1 in April (74% and 76% of the amino acid sequences were unique, respectively; Fig. 4, Figs. S6-S8). The least diverse sequences were from stations A4 and H4 in February (25% and 36% unique, respectively).

**Fig. S2.**
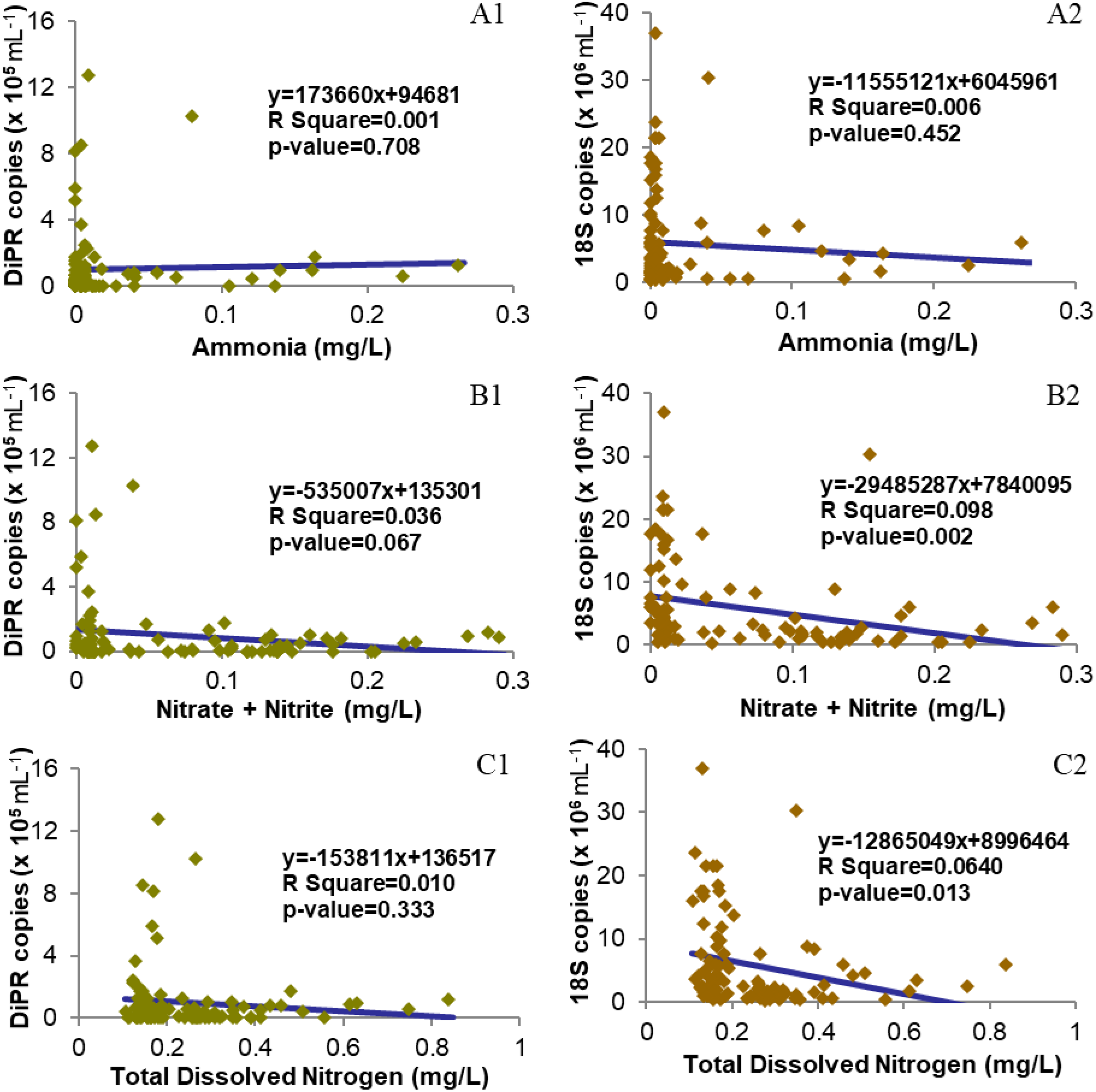
Scatter charts of gene copy numbers of dinoflagellate proton-pump rhodopsin (DiPR; 1) or phytoplankton 18S RNA (2) vs environmental factors with the best fit Line. A, Ammonia; B, Nitrate + Nitrite; C, Total Dissolved Nitrogen.

**Fig. S3.**
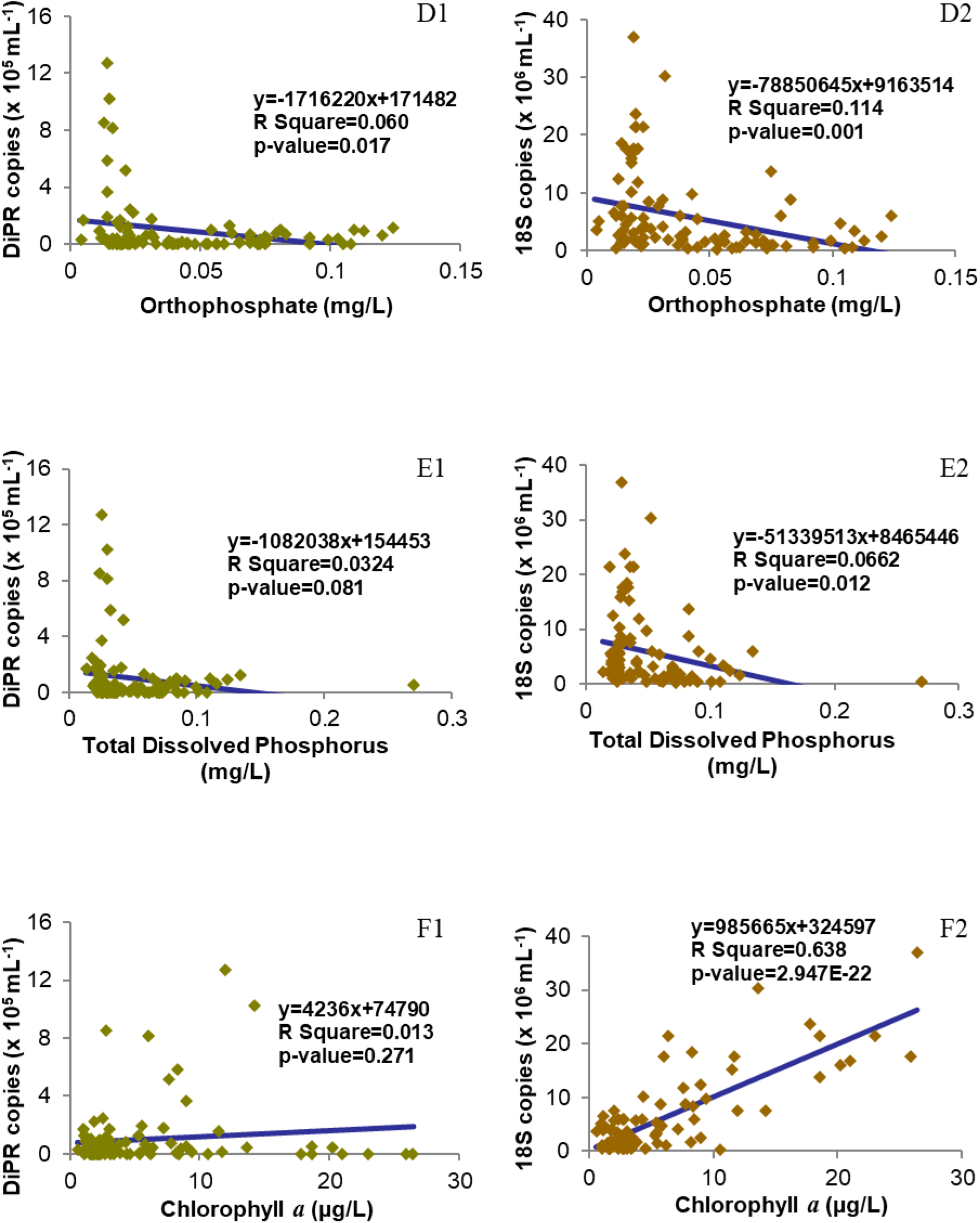
Scatter charts of gene copy numbers of dinoflagellate proton-pump rhodopsin (DiPR; 1) or phytoplankton 18S RNA (2) vs environmental factors with the best fit Line. D, Orthophosphate E, Total Dissolved Phosphorus; F, Chlorophyll *a.*

**Fig. S4.**
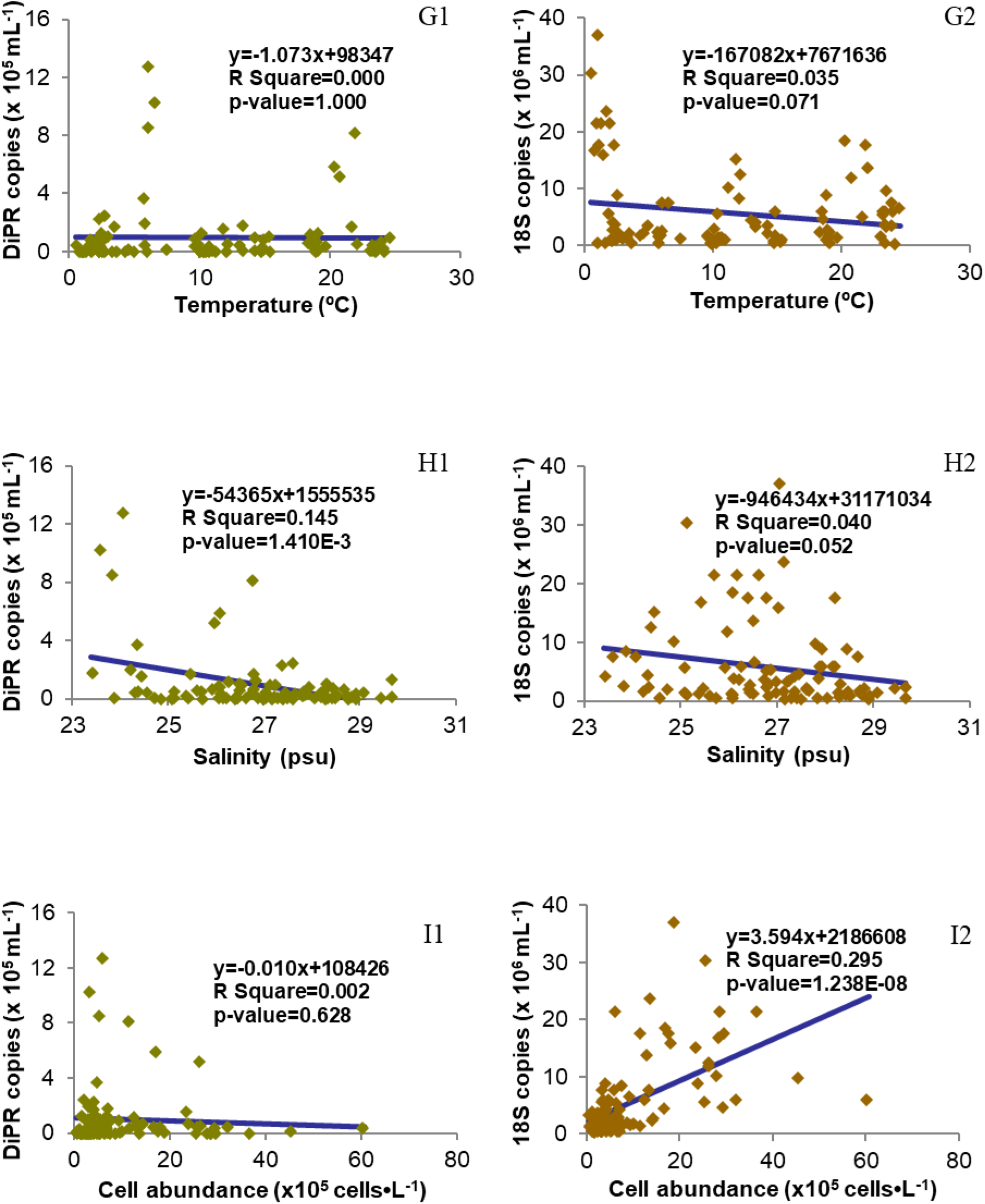
Scatter charts of gene copy numbers of dinoflagellate proton-pump rhodopsin (DiPR; 1) or phytoplankton 18S RNA (2) vs environmental factors (or cell abundance) with the best fit Line. G, Temperature; H, Salinity; I, Total phytoplankton cell abundance.

**Fig. 5.**
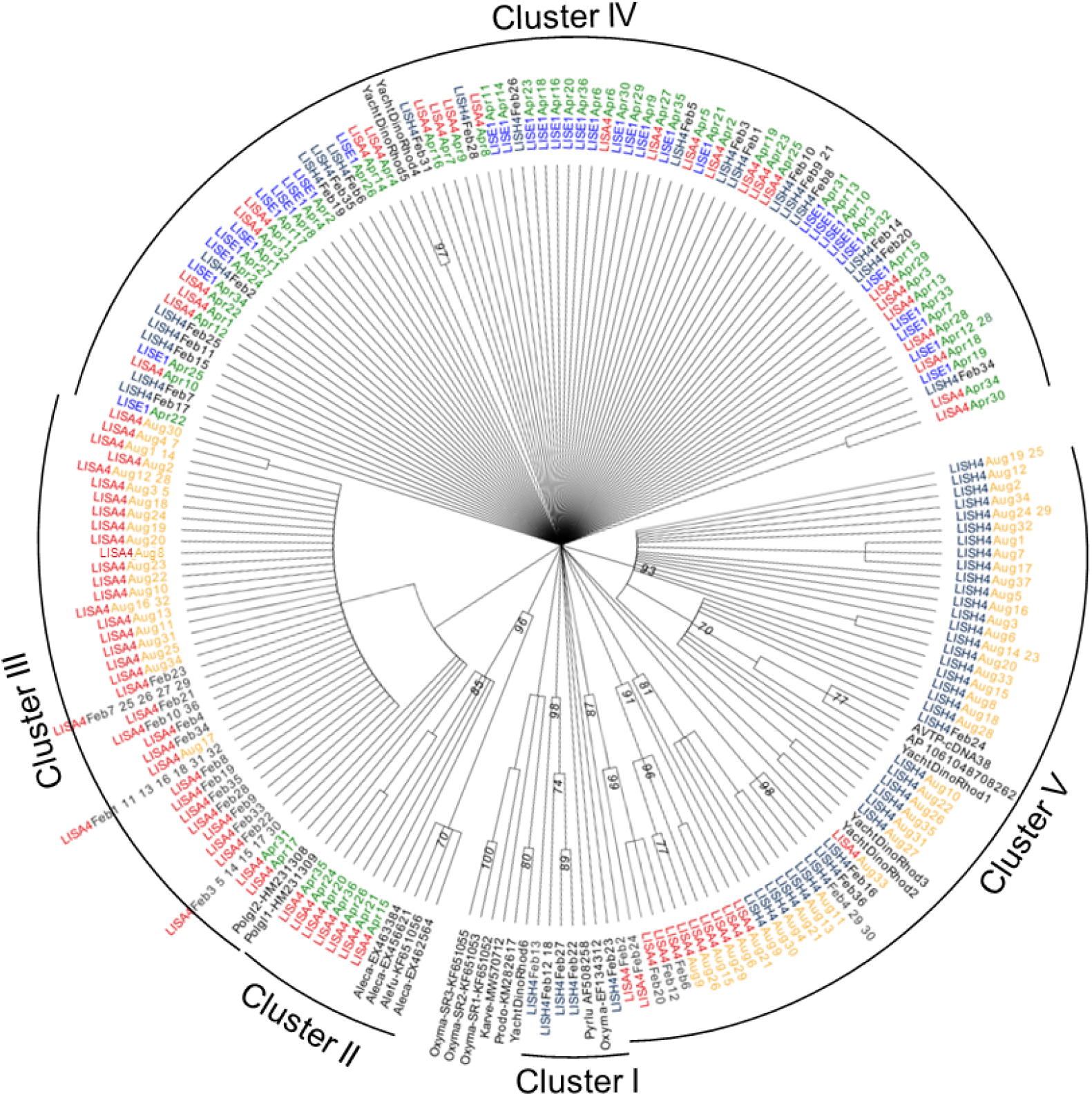
Maximum Likelihood tree of dinoflagellate rhodopsin genes obtained in this study. The representatives of the dinoflagellate rhodopsin genes reported in GenBank are included.

Phylogenetic analyses revealed that the sequences could be grouped into five clusters (Fig. 5). In Cluster I, there were six sequences (two identical), all of which were from the Station H4 February water sample. These sequences were closest to *O. marina* proton-pump rhodopsin.

Cluster II had eight sequences solely from water samples at Station A4 in April and were tightly grouped with *Alexandrium* PPR (Fig. S6).

Cluster III was composed of sequences solely from the Station A4 water samples in February (36 sequences), April (1) and August (26). These sequences were closely related to *Polarella glacialis* proton-pump rhodopsins.

Cluster IV contained sequences from all three tested stations in February and April, including A4 in April (26), E1 in April (34), and H4 in February (21). These sequences are grouped with the other two full-length dinoflagellate proton-pump rhodopsins obtained in this study. In this cluster, sequences from different stations in the same month or same station in different months could be clustered together, suggesting that some similar DiPPRs, and hence similar dinoflagellate taxa, exist throughout the Sound all year round.

Cluster V consisted of sequences mainly from the Station H4 August water sample (37), but some from H4 February (6) and A4 August water samples (7). These sequences were clustered together with the PPRs from unknown dinoflagellates we obtained in the previous study and three full-length PPRs we isolated from the water samples near our laboratory (Fig. 5; see next section for details).

### Full-length cDNAs of DiPPRs from the field

Six full-length DiPPR cDNAs from the water sample collected at our campus (YachtDinoRhod1 to 6) and two full-length rhodopsin cDNAs from *K. veneficum* were deposited to GenBank with Accession # MW570706-MW570713. YachtDinoRhod1 shared high nt and aa similarities with the two sequences we obtained previously from the water sample at a similar location (AVTP-cDNA38 GU553957 and AP_1061048708262 GU554267; Fig. 4, Figs. S6-S8; [8]). YachtDinoRhod2 and 3 are clustered with the sequences obtained from Station H4 in August. YachtDinoRhod4 and 5 are grouped with many sequences from water samples collected in all three stations (Fig. 5). YachtDinoRhod6, however, had a distinct sequence from the other five sequences, sharing some similarity with *P. minimum* rhodopsin yet with low bootstrap support.

## Discussion

Rhodopsin-based phototrophy is now recognized as an important mechanism of solar energy capture in the marine ecosystem [3,4,6,9,10]. By measuring *all-trans* retinal, chlorophyll-*a,* and bacteriochlorophyll-*a* abundance in the Mediterranean Sea and the Atlantic Ocean, Gómez-Consarnau and colleagues discovered that proton-pump proteorhodopsin-based phototrophy might contribute the same amount of light energy fixation as chlorophyll-*a* based phototrophy does, producing energy sufficient to support basal metabolism of bacteria in the surface ocean [7]. However, this method cannot distinguish whether the proton-pump proteorhodopsin-based phototrophy is from bacteria or picodinoflagellates or other picoeukaryotes. Using metatranscriptomics coupled with 18S rDNA tag sequencing method, Vader et al. [5] examined the expression of microbial proton-pumping rhodopsins and other genes in the mid-summer for function and composition of marine protists (size 0.45-10 μm) in the high-Arctic Billefjorden. No study, however, has been dedicated to understanding the seasonal variation of DiPPR gene abundance in the field. The present study provides the first documentation of the seasonal distribution pattern of DiPPR gene in the LIS estuary off the Atlantic Ocean.

### Dinoflagellate specific PPR primers

We designed primers at the conserved regions of known typical DiPPR (Table 2). Most of the reported DiPPRs, including those from *P. lunula* [15], *O. marina* [8,10,16,17], *P. glacialis* [8] and *A catenella* [10], belong to proton-pump rhodopsin from *r*-proteobacteria, and share >70% nt and >80% aa similarities. The more recently reported rhodopsin sequences from *Prorocentrum donghaiense* [18] and *Karlodinium veneficum* [19, this study], however, were not included in the alignment for primer designing. These sequences are very distinct, sharing <45% of aa similarity with typical DiPPR (Fig. 4). Likely, the DiPPR primers we designed did not amplify rhodopsin genes for *Prorocentrum* and *Karlodinium* species in the field even though both species occur in LIS [29,30]. When testing the DiPPR primers for both the cDNA and gDNA of the phytoplankton species listed in Table 1, we discovered that for some tested dinoflagellates, i.e., *Alexandrium pacificum*, *Pyrocystis lunula*, *P. noctiluca,* and *Symbiodinium microadriaticum*, only cDNA gave positive amplification. This might be due to the existence of intron(s) in the targeted PCR region, resulting in unsuccessful amplification of DiPPR from gDNA. No amplification appeared for any of the non-dinoflagellate algae tested, indicating these primers are dinoflagellate specific or that those other algae do not possess PPR. Also, these primer sets did not amplify any sensory type rhodopsins when cloned cDNA of *O. marina* sensory type rhodopsins [10] were used as templates. As such, the present study provides a typical DiPPR-specific primer set for future use in investigating the diversity and abundance of dinoflagellate PPR in natural environments.

### Spatial and seasonal changes of dinoflagellate PPR gene abundance in Long Island Sound

With the DNA extracted from a year’s worth of samples (95 in total), 18S rDNA qPCR successfully amplified 18S rDNA from each sample, and its copy numbers exhibited a similar dynamic pattern to the total phytoplankton cell abundance from microscopic counts (Figs. 3, 4, Figs. S1A, S1C, S4-I2), and corresponded well with chlorophyll *a* concentration in each sample (S3-F2). This verifies that the DNA quality and PCR conditions were reliable. From these DNA samples, qPCR results showed that the monthly gene abundance of DiPPR varied markedly among stations (Figs. 3, 4).

The temporal change of DiPPR abundance in LIS showed two major PPR abundance peaks, one in April (also high in March) and the other in July (Fig. 4A). The July peak could correspond to the high average dinoflagellate cell abundance in this month. However, in April, total dinoflagellate cell abundance was rather low at most of the stations, yet, DiPPR gene abundance was very high at stations A4, B3, and E1 (Fig. S1). As a result, when DiPPR gene abundance was normalized to the monthly average dinoflagellate cell number, only the April peak remained (Fig. 4B). When taking a closer look at the dinoflagellate species in March and April, we found that the dinoflagellate *Peridinium quinquecorne* was abundant at stations E1, F2, H4, I2, J2, and K2 in March, and A4, B3, C1, D3, E1 and F2 in April. We are not certain whether this dinoflagellate was the major contributor to the high DiPPR gene abundance during these months because we do not have this species in culture to confirm its gene sequence. On the other hand, the high dinoflagellate cell abundance in July was largely contributed by *Prorocentrum triestinum*, which most likely has similar rhodopsin with *P. minimum* and would not be detected with our DiPPR qPCR primers. This may explain why DiPPR abundance per dinoflagellate cell was low in July.

Spatially, DiPPR gene abundance showed a weak trend of being greater in Western Sound than in Central and Eastern Sound (Fig. 4C). However, when normalized to the monthly average dinoflagellate cell abundance, DiPPR gene abundance showed peaks at stations E1, J2 and K2 (Fig. 4D, Fig. S1). It is important to note that inclement weather prevented sampling at stations J2 and K2 for several months (4 months and 5 months, respectively) in 2010 (Table S1, Fig. S1), therefore abundance data could be biased for these stations. In addition, the peak at station E1 is primarily caused by the high DiPPR copy number in some species in April, possibly contributed by *P. quinquecorne*, and a generally low abundance of dinoflagellate cell number in this station.

We did not observe a clear correlation between DiPPR gene abundance and any particular environmental factor other than weak negative correlations with orthophosphate concentration and salinity (Figs. S2–S4). In April, when DiPPR gene copy number was the highest, the values of all the measured environmental factors (e.g., salinity, water temperature, chlorophyll *a*, concentration of ammonia, nitrate + nitrite, orthophosphate, total dissolved nitrogen and total dissolved phosphorus) were low in general. Similarly, in July when DiPPR gene copy number was also high, all of the environmental factors except water temperature were low. This result suggests that DiPPR gene abundance in LIS is not influenced by water temperature or nitrogen nutrient concentration. The observed negative correlation of DiPPR gene abundance with salinity is interesting, as it suggests that the DiPPR-harboring dinoflagellates might be slightly favored by lower salinities. In addition, the negative correlation of DiPPR abundance with phosphate is reminiscent of recent findings that PPR gene expression was up-regulated under phosphorus stressed condition in *P. donghaiense*, in which DiPPR is postulated to facilitate the dinoflagellate to endure or thrive under phosphate deficiency [18,20]. The functional association of DiPPR with phosphorus nutrition warrants further investigation in the future.

The seasonal variation in 18S rDNA gene abundances were similar to that of phytoplankton cell abundances and chlorophyll *a* concentrations (Figs. S1, S3) in LIS. The strong positive relationship between phytoplankton18S rDNA gene abundance and chlorophyll *a* concentration (R^2^ =0.638, p-value <0.01; Fig S3) indicates that our quantitative Real-Time PCR system worked properly, and the 18S rDNA abundance data are reliable proxies of phytoplankton cell abundance in a community. The weak negative correlations of 18S rDNA copy number with orthophosphate and nitrogen (other than ammonia) concentrations might be because the growth of phytoplankton consumed some of the nitrogen and phosphorus nutrients.

### DiPPR gene diversity in LIS

To explore the diversity of dinoflagellate PPR genes in LIS, we chose six DNA samples for cloning and sequencing of the dinoflagellate PPR genes, including three samples from A4 in February, April, and August, one sample from station E1 in April, and two samples from station H4 in February and August. These samples were chosen to get representatives to investigate the diversity of dinoflagellate rhodopsin by location in LIS (western vs. central) and by season (winter, spring, and summer). In addition, there was a peak of DiPPR abundance in April, and we intended to explore what kind of dinoflagellates they were.

The 209 sequences obtained could be grouped into five clusters, but none of them were identical to the few reported DiPPR genes (Fig. 4, Fig. S2); therefore, we cannot attribute them to specific dinoflagellate species. Cluster I contained six sequences solely from the Station H4 February water sample that was somewhat close to *O. marina* PPR, Cluster III were sequences from the water sample at Station A4 in April that were similar to *Alexandrium* PPR, while cluster II composed of sequences from Station A4 water samples in all the investigated months that were grouped with *P. glacialis* PPR. Cluster IV had sequences mostly from Station H4 in August, and some from H4 in February and A4 in August, while Cluster V contained sequences from all three tested stations in February and April but not in August. These results indicate that some dinoflagellates exist only at certain time of year at a specific location in the Sound, while others can be found at stations all year round. The functions and ecological implications of the DiPPR diversity pattern remain to be uncovered.

## Acknowledgments

We thank Matthew Lyman, Katie O’Brien-Clayton as well as the DEEP seasonal monitoring staff from the CT DEEP and the crew of Dempsey for collecting the samples and measuring the environmental variables from the LIS WQMP. This research was supported by the National Science Foundation’s EAGER grant #OCE-1212392 and the CT DEEP grant entitled “Identification of Phytoplankton collected from Long Island Sound”. All the authors declare no conflict of interest existing in this work.

## References

1. Falkowski PG. The role of phytoplankton photosynthesis in global biogeochemical cycles. Photosynth Res. 1994; 39: 235–258.

2. Field CB, Behrenfeld MJ, Randerson JT, Falkowski PG. Primary production of the biosphere:integrating terrestrial and oceanic components. Science. 1998; 281: 237–240.

3. Beja O, Aravind L, Koonin EV, Suzuki MT, Hadd A, Nguyen LP, et al. Bacterial Rhodopsin: Evidence for a New Type of Phototrophy in the Sea. Science. 2000; 289: 1902–1906.

4. Beja O, Spudich EN, Spudich JL, Leclerc M, DeLong EF. Proteorhodopsin phototrophy in the ocean. Nature. 2001; 411: 786–789.

5. Vader A, Laughinghouse IV HD, Griffiths C, Jakobsen KS, Gabrielsen TM. Proton-pumping rhodopsins are abundantly expressed by microbial eukaryotes in a high-Arctic fjord. Environ Microbiol. 2018; 20: 890–902.

6. Gómez-Consarnau L, González JM, Riedel T, Jaenicke S, Wagner-Döbler I, Sañudo-Wilhelmy SA, et al. Proteorhodopsin light-enhanced growth linked to vitamin-B1 acquisition in marine Flavobacteria. ISME J. 2016; 10: 1102–1112.

7. Gómez-Consarnau L, RavenNaomi JA, Levine NM, Cutter LS, Wang D, Seegers B, et al. Microbial rhodopsins are major contributors to the solar energy captured in the sea. Science Advances. 2019; 5(8): eaaw8855. DOI:10.1126/sciadv.aaw8855

8. Lin S, Zhang H, Zhuang Y, Tran B, Gill J. Spliced leader-based metatranscriptomic analyses lead to recognition of hidden genomic features in dinoflagellates. PNAS. 2010; 107: 20033–20038.

9. Ernst OP, Lodowski DT, Elstner M, Hegemann P, Brown LS, Kandori H. Microbial and animal rhodopsins:Structures, functions, and molecular mechanisms. Chem Rev. 2014; 114: 126–163.

10. Guo Z, Zhang H, Lin S. Light-Promoted Rhodopsin Expression and Starvation Survival in the Marine Dinoflagellate Oxyrrhis marina. PLoS ONE. 2014; 9(12): e114941. DOI:10.1371/journal.pone.0114941

11. Marchetti A, Catlett D, Hopkinson BM, Ellis K, Cassar N. Marine diatom proteorhodopsins and their potential role in coping with low iron availability. ISME J. 2015; 9: 2745–2748.

12. Sabehi G, Loy A, Jung K-H, Partha R, Spudich JL, Isaacson T, et al. New insights into metabolic properties of marine bacteria encoding proteorhodopsins. PLOS Biol. 2005; 3: e273.

13. Campbell BJ, Waidner, LA, Cottrell MT, Kirchman DL. Abundant proteorhodopsin genes in the North Atlantic Ocean. Environ Microbiol. 2008; 10: 99–109. doi:10.1111/j.1462-2920.2007.01436.x

14. Maresca JA, Miller KJ, Keffer JL, Sabanayagam CR, Campbell BJ. Distribution and Diversity of Rhodopsin-Producing Microbes in the Chesapeake Bay. Appl Environ Microbiol. 2018; 84(13): e00137–18.

15. Okamoto OK, Hastings JW. Novel dinoflagellate clock-related genes identified through microarray analysis. J Phycol. 2003; 39: 519–526.

16. Zhang H, Hou Y, Miranda L, Campbell DA, Sturm NR, Gaasterland T, et al. Spliced leader RNA trans-splicing in dinoflagellates. PNAS USA. 2007; 104: 4618–4623.

17. Slamovits CH, Okamoto N, Burri L, James ER, Keeling PJ. A bacterial proteorhodopsin proton pump in marine eukaryotes. Nature Communications. 2011; 183: doi:101038/ncomms1188

18. Shi X, Li L, Guo C, Lin X, Li M, Lin S. Rhodopsin gene expression regulated by the light dark cycle, light spectrum and light intensity in the dinoflagellate *Prorocentrum*. Front Microbiol. 2015; 6: 555.

19. Meng R, Zhou C, Zhu X, Huang H, Xu J, Luo Q, et al. Critical light-related gene expression varies in two different strains of the dinoflagellate *Karlodinium veneficum* in response to the light spectrum and light intensity. J Photochem Photobiol B Biol. 2019; 194: 76–83.

20. Zhang Y, Lin X, Shi X, Lin L-X, Luo H, Li L, et al. Metatranscriptomic signatures associated with regime shift from diatom dominance to a dinoflagellate bloom. Front Microbiol. 2019; 10: 590 Doi:103389/fmicb201900590.

21. Yu L, Zhang Y, Li M, Wang C, Lin X, Li L, Guo S, Lin S. Comparative metatranscriptomic profiling and microRNA sequencing to reveal active metabolic pathways associated with a dinoflagellate bloom. Sci Tot Environ. 2020; 699: 134323. https://doiorg/101016/jscitotenv2019134323

22. Zhang H, Lin S. Detection and quantification of *Pfiesteria piscicida* using mitochondrial cytochrome b gene sequence. Appl Environ Microbiol. 2002; 68: 989–994.

23. Zhang H, Lin S. Development of a *cob-18S* dual-gene Real-Time PCR assay for *Pfiesteria shumwayae* and quantification of this species in the natural environment. Appl Environ Microbiol. 2005; 71: 7053–7063.

24. Zhang H, Lin S. Status of mRNA editing and SL RNA trans-splicing groups *Oxyrrhis, Noctiluca, Heterocapsa*, and *Amphidinium* as basal lineages of dinoflagellates. J Phycol. 2008; 44: 703–711.

25. Finiguerra M, Avery DE, Dam HG. No evidence for induction or selection of mutant sodium channel expression in the copepod *Acartia husdsonica* challenged with the toxic dinoflagellate *Alexandrium fundyense*. Ecol Evol. 2014; 4(17): 3470–3481. DOI:10.1002/ece3.1197

26. Hasegawa M., Kishino H., and Yano T. Dating the human-ape split by a molecular clock of mitochondrial DNA. J Mol Evol, 1985; 22:160–174.

27. Felsenstein J. Confidence limits on phylogenies: An approach using the bootstrap. Evol. 1985; 39:783–791.

28. Kumar S., Stecher G., Li M., Knyaz C., and Tamura K. MEGA X: Molecular Evolutionary Genetics Analysis across computing platforms. Mol Biol Evol. 2018; 35:1547–1549.

29. Lonsdale DJ, Greenfield DI, Hillebrand EM, Nuzzi R, Taylor GT. Contrasting microplanktonic composition and food web structure in two coastal embayments (Long Island, NY, USA). J Plankton Res. 2006; 28 (10): 891–905.

30. Zhang H, Litaker W, Vandersea MW, Tester P, Lin S. Geographic distribution of *Karlodinium veneficum* in the US east coast as detected by ITS-ferredoxin Real-Time PCR assay. J Plankton Res. 2008; 30: 905-922

31. Shi X, Lin X, Li L, Li M, Palenik B, Lin S. Transcriptomic and microRNAomic profiling reveals multi-faceted mechanisms to cope with phosphate stress in a dinoflagellate ISME J. 2017; 11: 2209–2218.

